# “A novel paradigm for optimal mass feature peak picking in large scale LC-MS datasets using the ‘isopair’: isoLock, autoCredential and anovAlign”

**DOI:** 10.1101/2021.12.05.471237

**Authors:** Allen Hubbard, Louis Connelly, Shrikaar Kambhampati, Brad Evans, Ivan Baxter

## Abstract

Untargeted metabolomics enables direct quantification of metabolites without apriori knowledge of their identity. Liquid chromatography mass spectrometry (LC-MS), a popular method to implement untargeted metabolomics, identifies metabolites via combined mass/charge (m/z) and retention time as mass features. Improvements in the sensitivity of mass spectrometers has increased the complexity of data produced, leading to computational obstacles. One outstanding challenge is calling metabolite mass feature peaks rapidly and accurately in large LC-MS datasets (dozens to thousands of samples) in the presence of measurement and other noise. While existing algorithms are useful, they have limitations that become pronounced at scale and lead to false positive metabolite predictions as well as signal dropouts. To overcome some of these shortcomings, biochemists have developed hybrid computational and carbon labeling techniques, such as credentialing. Credentialing can validate metabolite signals, but is laborious and its applicability is limited. We have developed a suite of three computational tools to overcome the challenges of unreliable algorithms and inefficient validation protocols: isolock, autoCredential and anovAlign. Isolock uses isopairs, or metabolite-istopologue pairs, to calculate and correct for mass drift noise across LC-MS runs. autoCredential leverages statistical features of LC-MS data to amplify naturally present 13C isotopologues and validate metabolites through isopairs. This obviates the need to artificially introduce carbon labeling. anovAlign, an anova-derived algorithm, is used to align retention time windows across samples to accurately delineate retention time windows for mass features. Using a large published clinical dataset as well as a plant dataset with biological replicates across time, genotype and treatment, we demonstrate that this suite of tools is more sensitive and reproducible than both an open source metabolomics pipelines, XCMS, and the commercial software progenesis QI. This software suite opens a new era for enhanced accuracy and increased throughput for untargeted metabolomics.

## Introduction

Accurately identifying and quantifying metabolites in an untargeted fashion is an important goal for the medical and life sciences. If achieved, it will improve the ability to characterize biochemistry in disease and health states, improve establishment of better clinical biomarkers (Clish, 2015) as well as enable better understanding of biochemistry in organisms across all kingdoms. The orbitrap (Zubarov and Makarov, 2013) and other high-resolution machines, which offer mass resolution at the single digit parts per million (ppm) error range, now produce larger and more complex datasets. However, detection of meaningful signals still remains challenging because of the high degree of random noise incurred by these sensitive instruments, as well as the prevalence of contaminants such as salts. Novel informatics techniques are needed to increase the signal to noise ratio in the LC-MS data in order to accurately identify mass feature signals of metabolites. Despite decades of method development, LC-MS driven untargeted metabolomics faces numerous challenges which have only been partially solved (Gertsman and Barshop, 2018). In LC-MS, metabolites are identified as mass features, or regions of signal which occur at a unique point in the space of retention time and mass. The mass and retention time, are both stored in vendor specific binary files which are often converted to open source text formats (Martens *et al*., 2011). Unfortunately, these files are large and unwieldy, and contain abundant noise produced by the sensitive machinery. When existing algorithms analyze this data, the noise frequently causes peak splitting and other detection issues. The problems of distinguishing signal from noise are amplified when large numbers of samples are used (dozens to hundreds of samples) because a compound must be reproducibly tracked in each sample in order to be reliably detected in the dataset.

There are three major sources of noise that limit the ability of existing informatics approaches to reproducibly detect compounds across samples. Two of these sources of noise, mass drift between runs and across samples, relate to the mass spectrometry aspect of LC-MS. The third source of noise, retention time drift, derives from the high pressures needed to be maintained to ensure consistent elution from the chromatography column, and transient fluctuations.

### Mass Drift Decreases Resolution

Even with high resolution instruments, maintaining a mass resolution within intended ranges is difficult (Gorshkov *et al*., 2011). In order to resolve metabolites, mass spectrometry must be both accurate, meaning an ion of a certain mass will be detected at that mass using the instrumentation, and precise, meaning that repeated measurement of the given mass closely agree with one another across time (Brenton *et al*., 2010). Effective and frequent calibration is required to maintain suitable accuracy and precision. Initial calibration ensures that the machine accurately resolves a target ion to within an acceptable range of error, such as +/- 5 parts per million (ppm). To put this sensitivity in perspective, effective calibration at 5 ppm accuracy would ensure that all signals associated with a compounds of mass 100 Da will fall within 99.9995 Da to 100.0005 Da. Owing to practical limitations, high resolution mass spectrometers are usually calibrated once a week. A typical machine may then drift up to 1-3 ppm a day from its point of calibration, causing the precision to degrade (citation). Over time, this drift becomes substantial, effectively “smearing” mass signals between calibrations and reducing resolution. Frequent calibration is an imperfect solution to mass drift. This is because the accuracy of a single calibration attempt, while generally within 2-3 ppm (Hecht *et al*., 2019), will depend on uncontrollable external factors such as fluctuations in temperature, humidity or other stochastic variables. This results in a subtle batch effect, as each week’s calibration may disagree with one another by up to several ppm, causing decay of resolution. Taken together, variability in calibration efficacy and the degradation of precision between calibrations, are serious problems that reduce the resolution of high resolution mass spectrometers when datasets of hundreds to thousands of samples which must be run on timescales of weeks to months are being analyzed. Using spike-ins as internal standards is not an ideal solution to capture this variation, as the set of standards will be unlikely to match the diversity of compounds detect by untargeted metabolomics, and thus can introduce measurement bias. Additionally, these standards will be prone to their own peak picking concerns which would interfere with the ability to detect drift.

### Stochastic Noise Signals Interfere with Mass Identification

In addition to the smearing of signals from mass drift, the production of stochastic noise signals makes it difficult to resolve the masses of true mass features from noise (Du *et 2008*). Owing to the extreme sensitivity of high-resolution mass spectrometers, spurious m/z signal intensities are produced stochastically (Kaur *et al*., 2006). These signals are distributed randomly across masses, and their intensities range between 0 to 10^5 counts, overlapping with the abundance of many metabolites. The signal-to-noise ratio in single samples is frequently too low to delineate metabolite masses from noise, causing a proliferation of dropouts as the number of samples in a dataset increases. The high prevalence of missing values due to technical reasons limits the detection of important biology and has required statistical solutions (Wei *et al*., 2018).

### Chromatography Performance Is limited by Retention Time Drift

Once the mass of a metabolite is identified, the compound’s retention window in a chromatography column (such as HILIC or RPLC for polar and nonpolar compounds, respectively) must also be determined in order to define metabolite as a mass feature. Using retention time as an additional dimension of measurement enables separation of isomers (Pan *et al*., 2005). Consistent retention times can be difficult to maintain across multiple samples due to the high required pressures and the impact of transient temperature and pressure gradients (Asberg *et al*., 2017). Shifts in retention time can also cause false peak-splitting events and dropouts in large datasets.

### Limitations of Existing Methods to Cope with Noise

Existing peak-picking algorithms, such as centWave, identify metabolites by extracting signals concentrated in a unique window of mass and retention time. These are called regions of interest (ROI) (Tautenhahn *et al*, 2008). centWave attempts to overcome noise in both domains (mass drift or retention time drift) by using a 2-dimensional binning heuristic to define ROI’s, with the parameters of this binning procedure allowing variability in mass and retention time across samples. However, this paradigm creates serious trade-offs between sensitivity and specificity when defining ROI’s (Figure 2). If noise thresholds are set too stringently, the number of metabolites detected plummets. Conversely, if thresholds are too relaxed, noise regions are turned into false positive metabolite predictions. The strategy identifying metabolites as ROI’s through the 2-dimensional bins compounds uncertainty in both the mass and retention time domain and forces unacceptable tradeoffs between sensitivity and specificity, regardless of how parameters are determined. This causes both a high number of dropouts and false positives in large datasets. While companion software (such as IPO) can be used to tune parameters for these algorithms, performance gains will be limited by the drawbacks of ROI based algorithms. For example, XCMS, an open source software which implements centWave, finds ROI’s in each file individually and then attempts to align predictions across multiple samples (Alboniga *et al*., 2020). This strategy of analyzing individual files separately can propagate errors, even when optimize parameters are used. A new paradigm, which rigorously models and fully accounts for noise in LC-MS data, addressing each source independently, is needed to define metabolite mass features.

**Figure 1.**
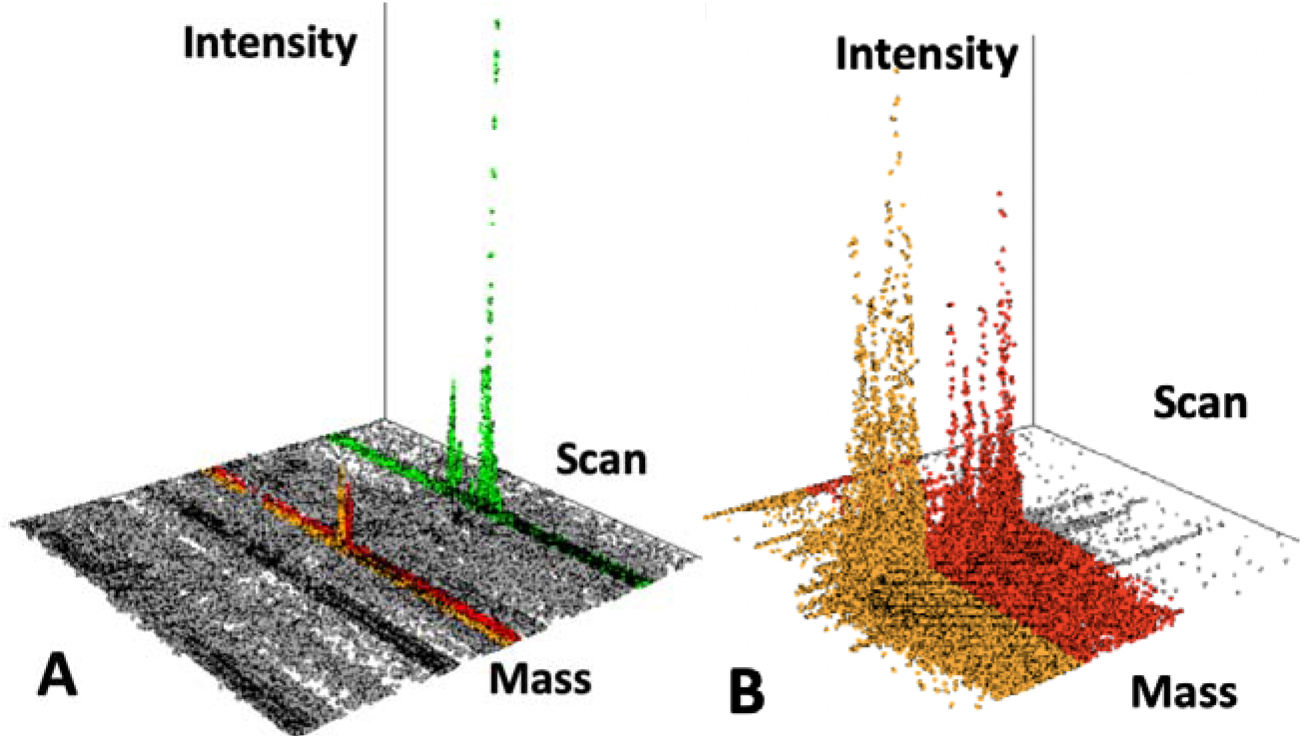
**(A & B):** Picking peaks in LC-MS data is challenging. A. Shown is a sliver of a chromatogram within +/- 200 ppm of the metabolite citrulline’s peak (176.1034 Da, red and orange regions). This represents only a fraction of fraction of a percent of the total data in an LC-MS run with a typical (m/z) range of 80-1000 Da. Signals from different metabolite mass features (colored) must be distinguished from noise (grey), as well as each other. Peak picking is even more complicated when hundreds of samples are pooled together, as mass and retention time drift cause serious issues for existing signal processing algorithms. This noise causes peak splitting in many algorithms (Fig 1B) (red and orange regions are actually a single peak, citrulline)

**Figure 2.**
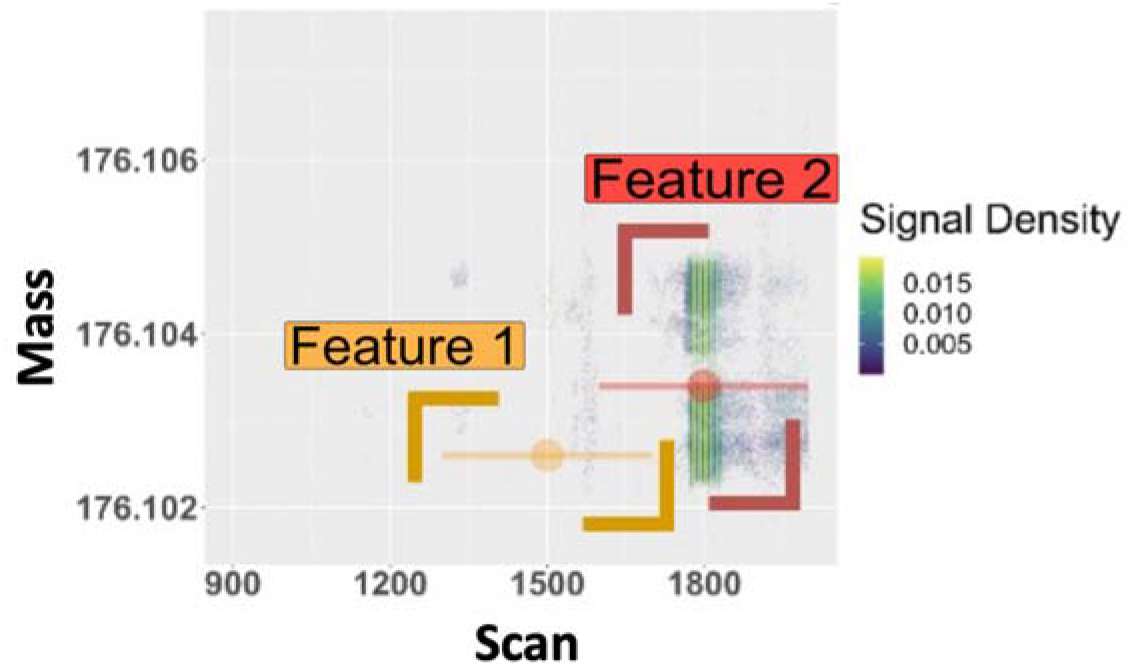
Existing algorithms such as centWave essentially view the 3-Dimensional mass feature peaks as 2D ion heatmaps, in which 2 dimensional sliding windows are fit around plausible regions of metabolite signals regions of interest (ROI’s). While powerful, this imperfect approach can split signals (Citrulline is split into two features) as noise accumulates across samples as well as miss true metabolites if filtering parameters are too stringent. Mass feature boundaries depicted in Figures 1 & 2, were determined using Progenesis QI on a previously published, 600 sample human microbiome dataset (Lloyd-Price *et al*., 2019).

**Figure 3:**
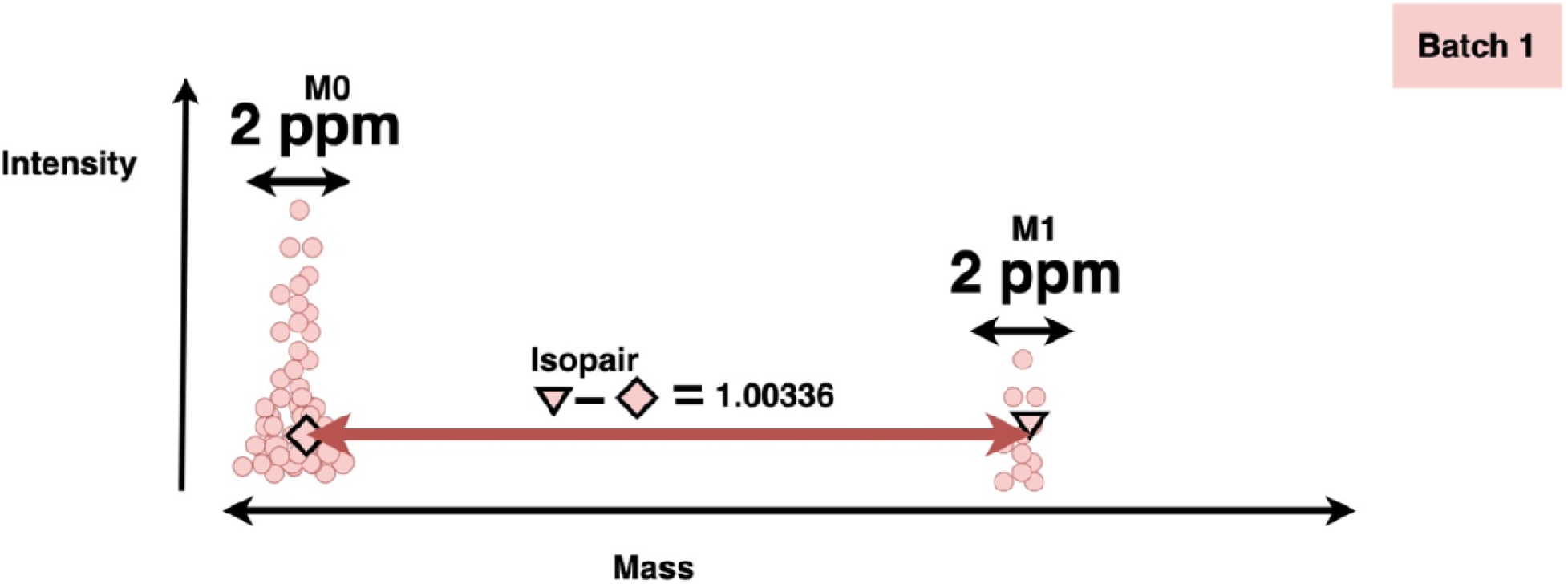
isopairs are signals separated by a mass shift corresponding to the approximate mass difference between 12C and 13C. This mass can now be determined to multiple decimal points using HRMS. Spread between peaks is not to scale.

### Big-Data oriented Solutions

One of the factors limiting approaches to handling metabolomics datasets was the large file sizes associated with each run. Increased availability of large-scale computing resources has made it possible to design novel peak picking algorithms which can effectively handle the multifaceted sources of LC-MS noise. We propose a novel software suite (isolock, autoCredential and anovAlign, from here on called the IAA suite) that is capable of analyzing large scale LC-MS datasets (hundreds to thousands of files) in a single analysis, thereby exploiting statistical features of the data to distinguish signal from noise in ways that are only evident when LC-MS files are pooled. This software suite uses 3 novel algorithms to first correct for mass drift (isoLock), then identifies masses most likely to belong to true metabolites (autoCredential), and, finally, correct for retention time drift (anovAlign) to refine the signals associated with these masses into high confidence mass features.

## RESULTS

### Growing the signal to noise ratio: Merged mass spectra and isopairs

When multiple samples are merged together into single mass spectra, there is an extreme amplification of signals associated with both metabolites (M0’s) and their substantially less abundant isotopologues (M1’ s). This amplification of isotopologues makes it possible to correspond mass-intensity signals of a metabolite with the signal of its isotopologue. These pairs of corresponding signals between a metabolite M0 and M1 will be equivalent to the mass gain derived from a 13C atom (1.00336 Da), which is resolvable using HRMS. These pairs of mass-intensity signals between metabolite M0’s and M1’s are known as isopairs. Because the masses determined by HRMS are accurate to several decimal places, and isopairs require two signals spaced at the exact mass of a 13C atom, the probability of two noise signals belonging to an isopair is extremely low.

### Isolock uses isopairs to correct for mass drift

Although signal to noise ratios increase when mass spectra are pooled together, mass drift interferes with successful alignment of signals in a merged spectra. When spectra from large batches are pooled together, mass signals can spread well outside the intended limits of high resolution mass spectrometers. Fig 4A illustrates an example of mass drift across a dataset of hundreds of samples.

**4A:**
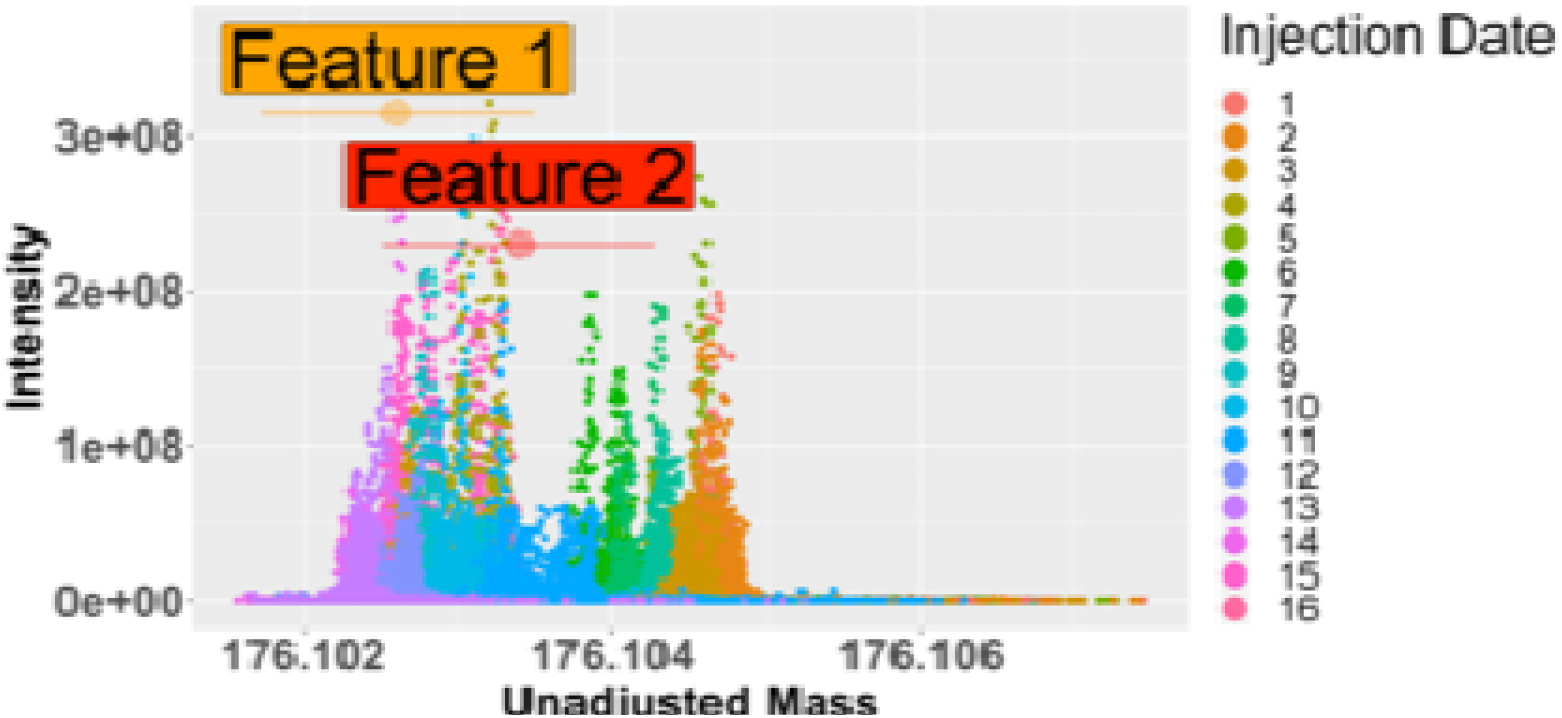
Citrulline, an abundant metabolite, demonstrates substantial mass drift across a set of 600 human fecal microbiome samples analyzed in batches across multiple days. This causes peak splitting when analyzed by existing algorithms.

Isopairs, however, can be used to capture the effect of drift across the mass spectra so that it can be removed (Figure 5). When mass drift is present, the separation between isopairs will no longer be equal to the mass gain of 13C atom, but 1.00336 + the mass drift (in ppm). The correct amount of mass drift between sets of samples can be determined by first selecting a sample early in the injection queue as a reference. The number of isopairs generated (across all masses) by the mass gain of a 13C atom + plausible mass drift values (+/- 20 ppm) can be calculated (Figure 6). Isolock determines the optimal mass drift value as that which maximizes isopairs and all mass values can then be adjusted by this value. After alignment of spectra via isoLock, spectra from hundreds to thousands of samples can be pooled into a single merged mass spectra in which signal to noise ratios are dramatically enhanced (Figure 4B).

**4B:**
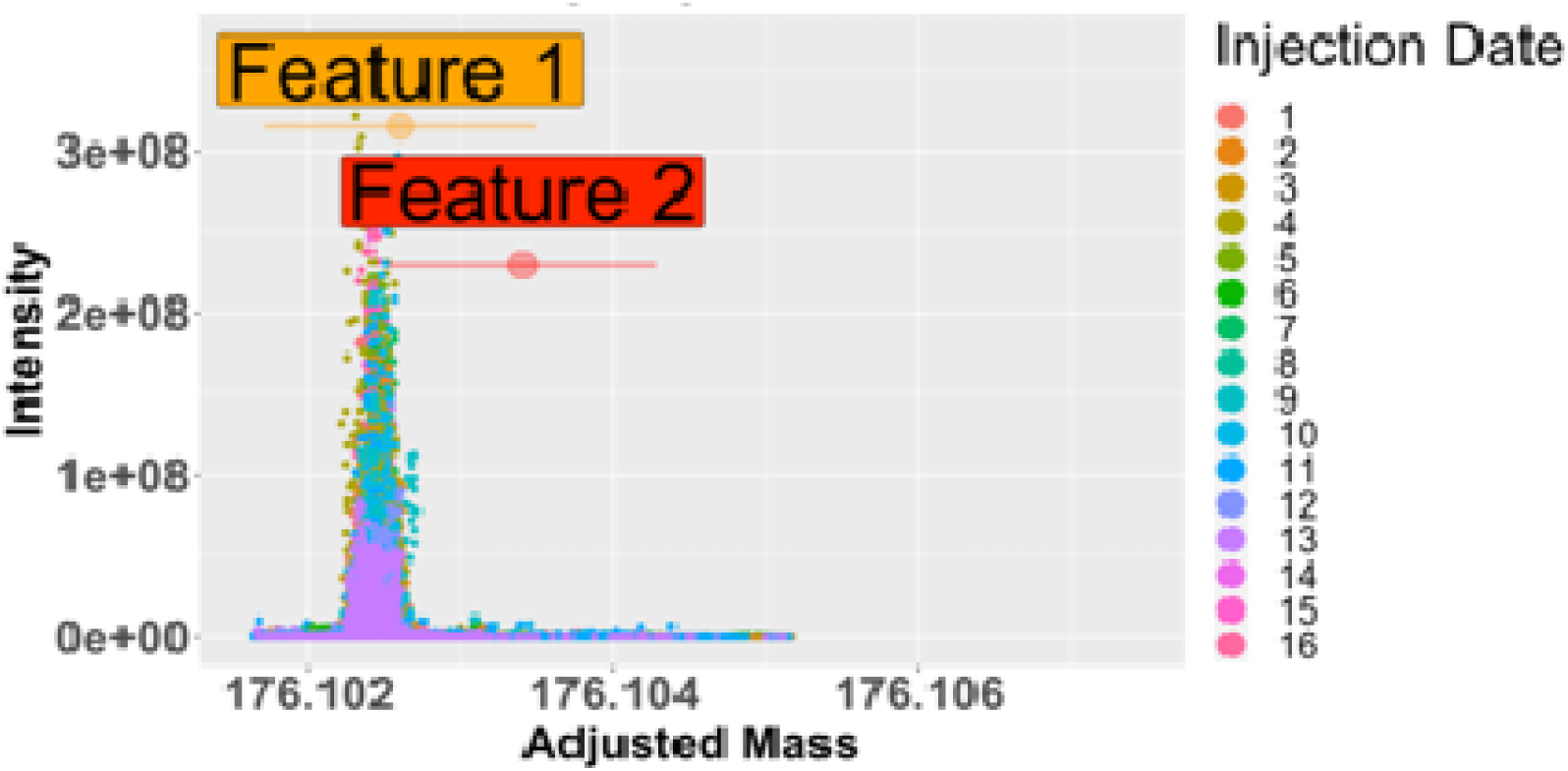
After correction for drift using the algorithm isoLock, the data is resolvable as a single peak.

**Figure 5:** Isopairs can quantify deviations in the measured mass from true mass.

**Figure 6:**
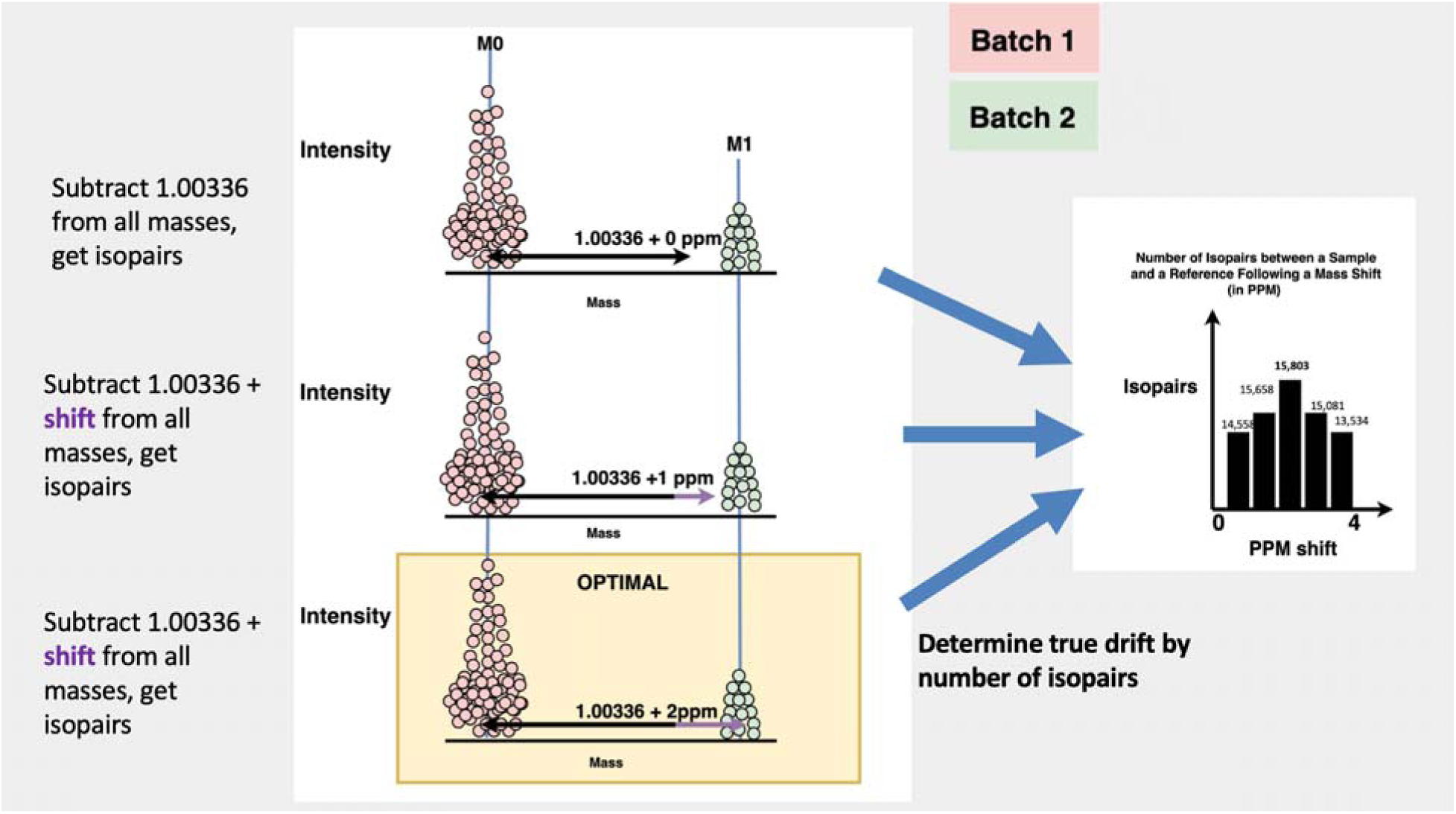
The determination by isoLock of a 2 ppm mass drift in a query sample relative to reference (via isopoairs) is shown. Isopairs are calculated in a fashion such that the M0 will belong to a reference file, and the M1 component will belong to a query file, with the separation between isopairs being allowed to vary across plausible mass shifts (+ the mass gain of 13C atom). The correct shift (2 ppm) is that which produces the highest number of isopairs.

### autoCredential uses background distribution and isopairs to identify high priority massintensities on a merged mass spectra

In a typical large-scale LC-MS metabolomics experiment, the combined mass spectra of hundreds of samples may contain billions of signals. However, these correspond to only thousands (at most) of metabolites (Mahieu *et al*., 2014). Thus, the vast majority of mass-intensity signals are noise. While the sheer number of noise signals is daunting, the statistical power resulting from a merged mass spectra enables effective filtering of metabolite masses from noise. The signal to noise ratio of a merged mass spectra is maximized by correcting for drift using isolock prior to merging spectra, as signals hyperconcentrate around the mass of valid mass features (Figure 7). While the vast majority of signals in a merged mass spectra are noise by raw number, they are spread out across the many millions of possible masses detectable by a high resolution instrument. Thus, their information density is low.

**Figure 7:**
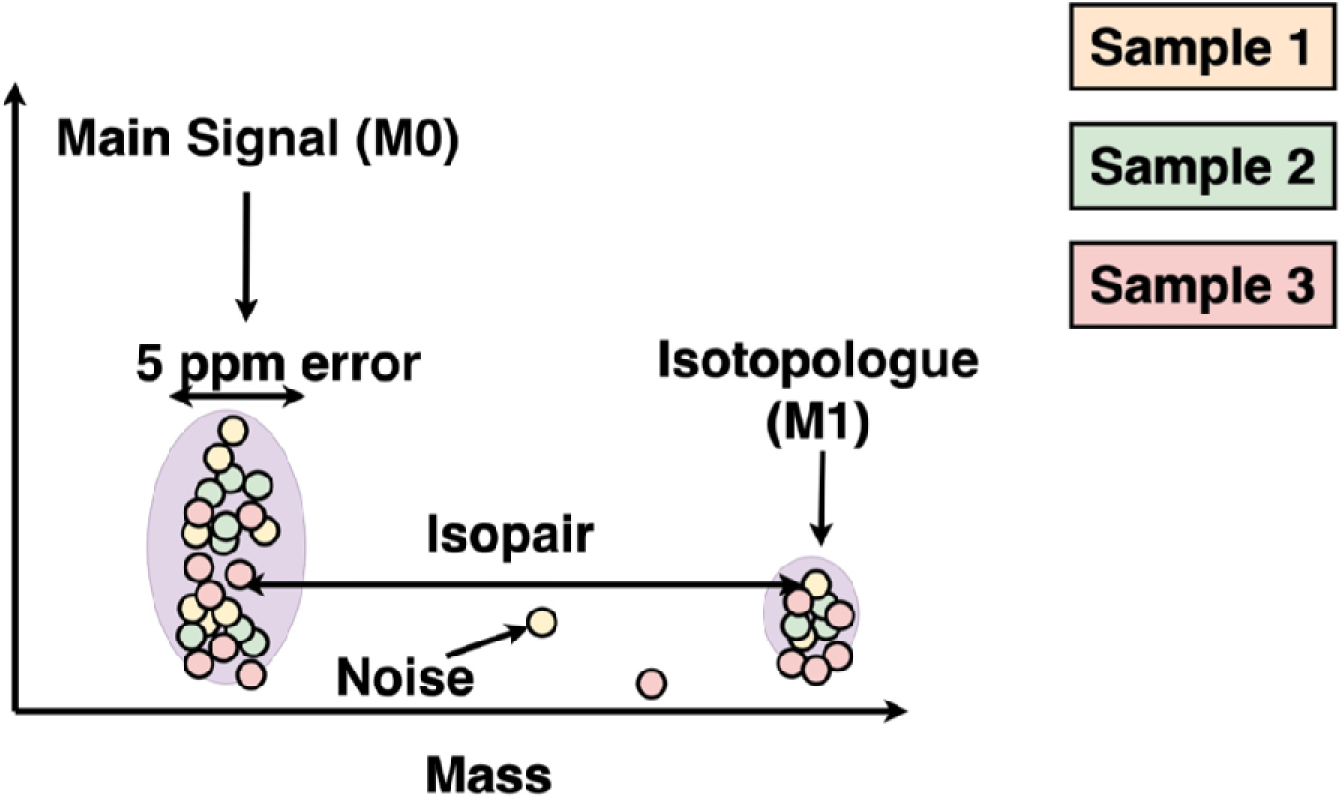
Signals around a metabolite (M0) and its isotopologue (M1) **increase** as samples are pooled together in a merged mass spectra. Because noise signals are random, noise regions do not show the dramatic growth in signal as samples are pooled.

**Figure 8:**
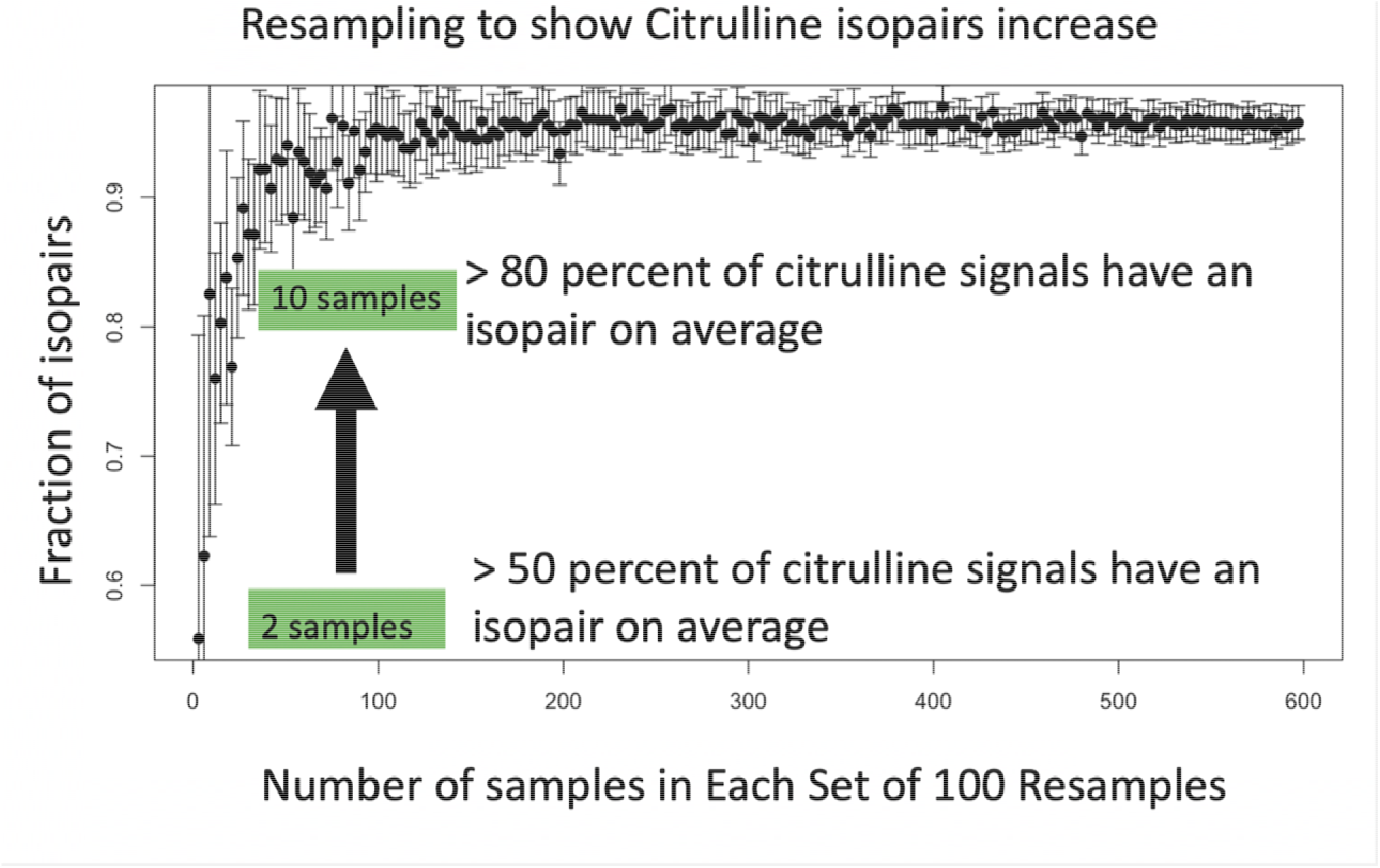
In the test dataset, we further checked to see if the masses were present in the majority of samples (>300), finding 80% of the masses met that criteria. To ascertain how many samples were required for autoCredential to effectively find isopairs for compounds, we resampled random combinations of the 600 runs of size 3-600 and tracked how many of our high-confidence metabolites had isopairs in each resample. We show that, on average, half the signals have isopairs with only two samples pooled into a merge mass spectra and that this fraction increases to eighty percent by the time 10 samples included in the merged mass spectra.

Dividing a merged mass spectra into .0001 Da bins (well within the idealized 1 ppm level of resolution for all relevant masses) and resampling at random, demonstrated that a single .0001 Da bin rarely contains more than several dozen noise signals even when hundreds of files are merged (Figure S1). Mass bins of this same size (.0001 Da) associated with either a metabolite or even its related and less abundant species (isotopes and adducts), however, will contain hundreds to thousands of signals. Thus, purely noise regions of a merged mass spectra can be pruned by removing any .0001 Da bin with less than several dozen signals. In our test set of 600 human fecal metabolome samples, this reduced >2 Billion masses to ~1 million masses. The merged mass spectra, however, still contains many artifactual signals, such as those belonging to inorganic salts. These can be removed by keeping only mass signals which are members of isopairs. Even if a singular region escapes denoising, the probability of detecting noise as a peak is especially low on a denoised mass spectra, as doing so requires two noise signals to occur at the exact interval equivalent to mass gain of a 13C atom is low.

Calculating isopairs reduced the number of plausible masses ~another order of magnitude. As highly abundant metabolites will spawn multiple distinct isopairs, collapsing isopairs within 1ppm yields a realistic number of metabolite masses (several dozens of thousands).

### anovAlign: Accounting for retention time drift to identify mass features

Although isolock corrects for mass drift, and autoCredential ensures that masses belong to metabolites (as opposed to inorganic salts - which would lack an isopair), retention time drift still makes it difficult to define the bounds of a mass feature. Once masses are accurately identified, chromatogram slices around a mass can be effectively extracted on a mass-specific basis for compound specific modeling. anovAlign uses signal to noise thresholds and identification of localized regions of scans whose signals are correlated to identify the likely region chromatographic peak of the signal. Once identified, the drift in each sample from the peak of a reference sample can be modeled as the translation of a gaussian, via anova (Figure 9). This is accomplished by running a t-test on the array of scan numbers for all samples compared to a reference sample (chosen as that with the highest signal). If the p-value is less than .05, the difference between the two means is subtracted from the query file. A signal-to-noise filter is rerun on this drift adjusted chromatogram slice to identify the refined mass feature bounds.

**Figure 9:**
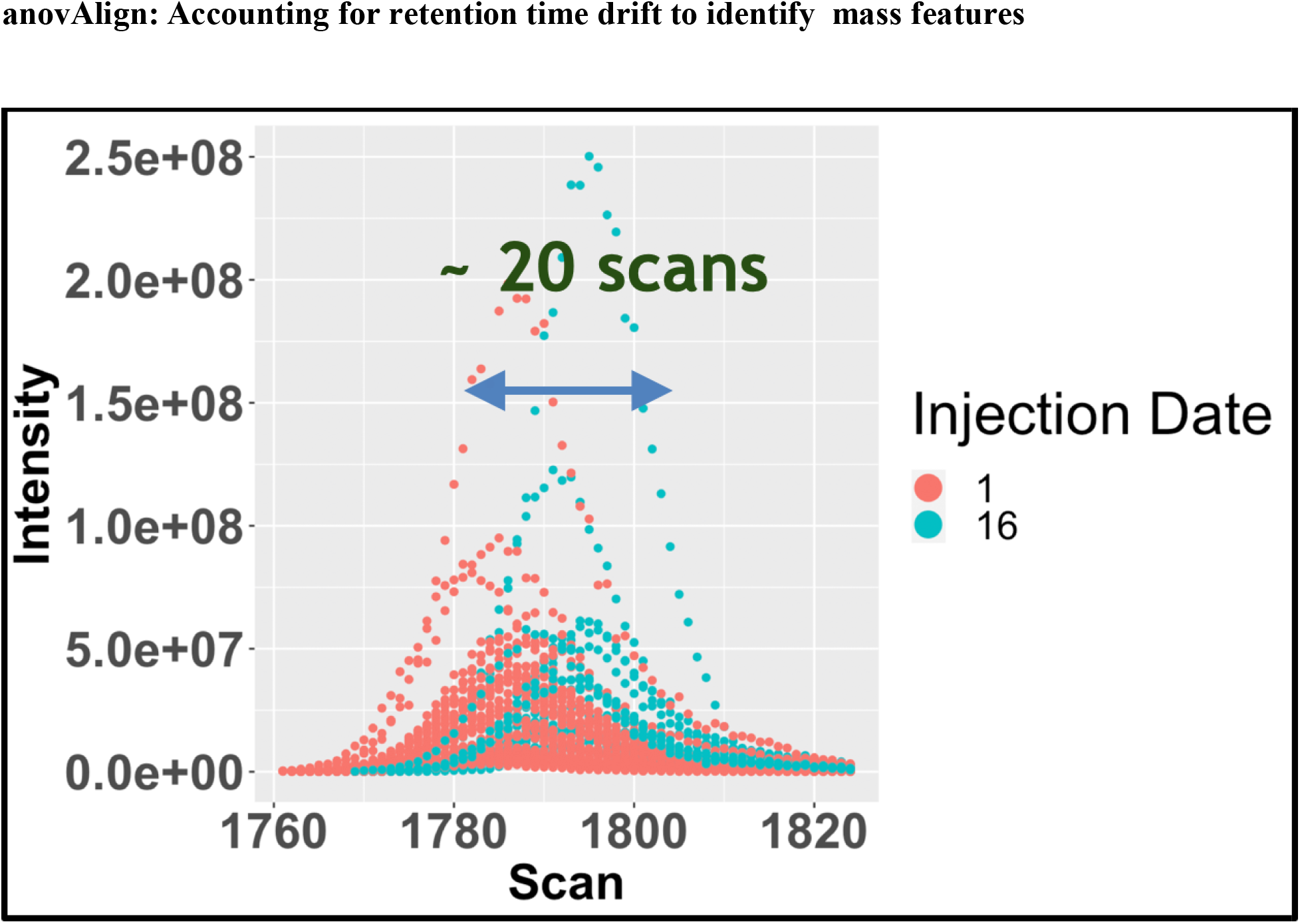
_The retention time of citrulline shows visible drift across the first batch (injection date 1) and the final batch (injection date 16).

Examining the prediction for citrulline mass feature using our suite (isolock, autoCredential, anovAlign) makes it clear that the paradigm of data pre-polishing (isoLock) followed by validation of masses via isopairs (autoCredential) and, finally, delineation and cleaning of the retention time bounds via anovAlign (Figures 10a and 10b). Considering the signal of a mass feature, such as citrulline, in isolation effectively captures the sequential progression of data through our pipeline. Much of the noise likely responsible for mass feature splitting in Progenesis QI prediction (Figure 10) resides in the mass domain. Once this is resolved, autoredential validates the correct mass using isopairs and the complete signal effectively falls out, allowing anovAlign to define the final mass feature boundaries (Figure 10b). This combined workflow of polishing **AND** feature selection effectively takes care of the “noisy data in, noisy data out” conundrum of traditional LC-MS software pipelines. While citrulline provides a good proof of concept as single signal in a large dataset, it is important to view the global performance of these algorithms in large, complex datasets.

**Figure 10 a:**
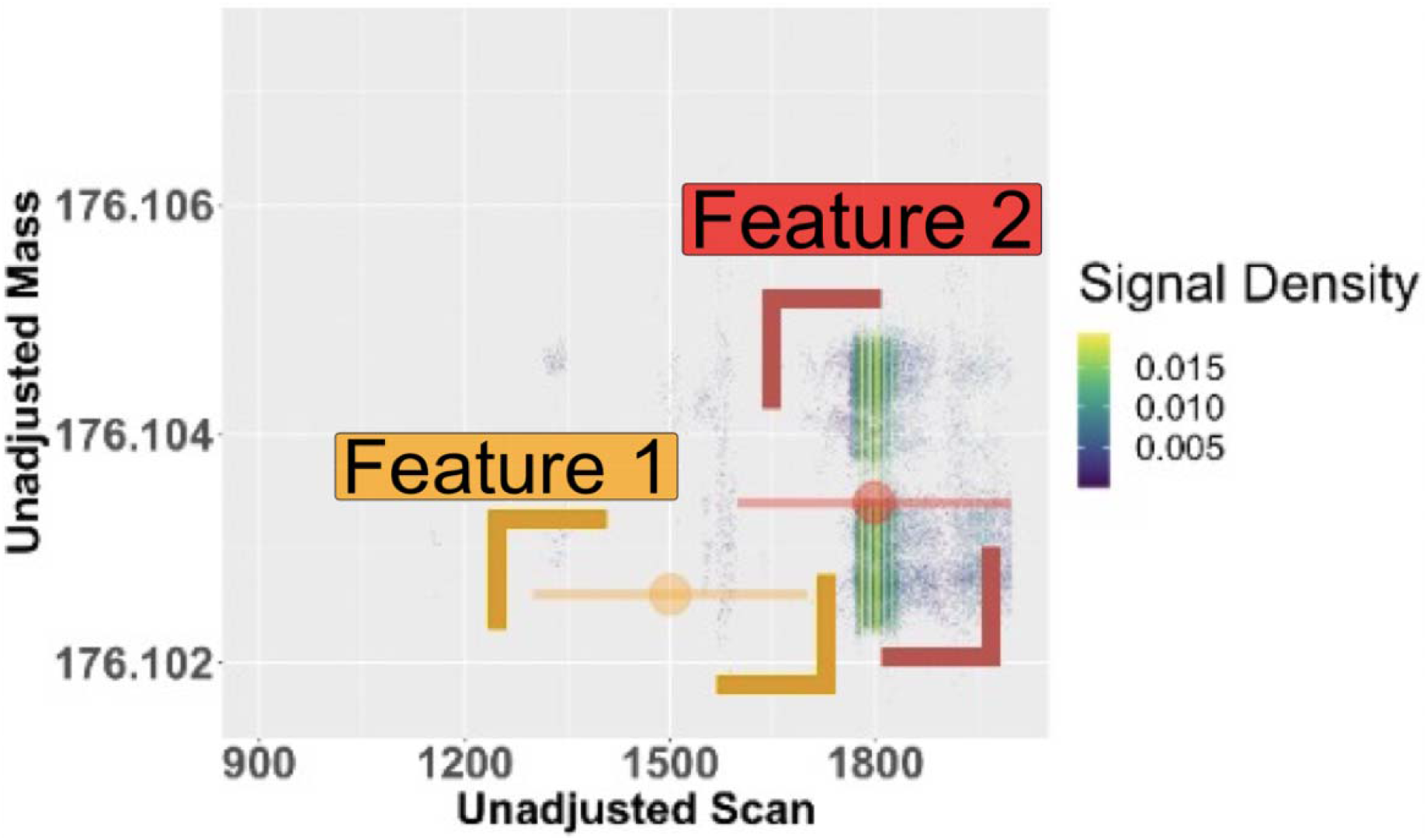
Before application of our pipeline (isoLock, autoCredential and anovAlign) the diffuse signal around a metabolite mass (such as citrulline) has substantial mass and RT drift which makes it difficult to call the mass feature correctly. This noise causes splitting in mass feature prediction using commercial software (the bounds determined by Progenesis QI are shown).

**Figure 10 b:**
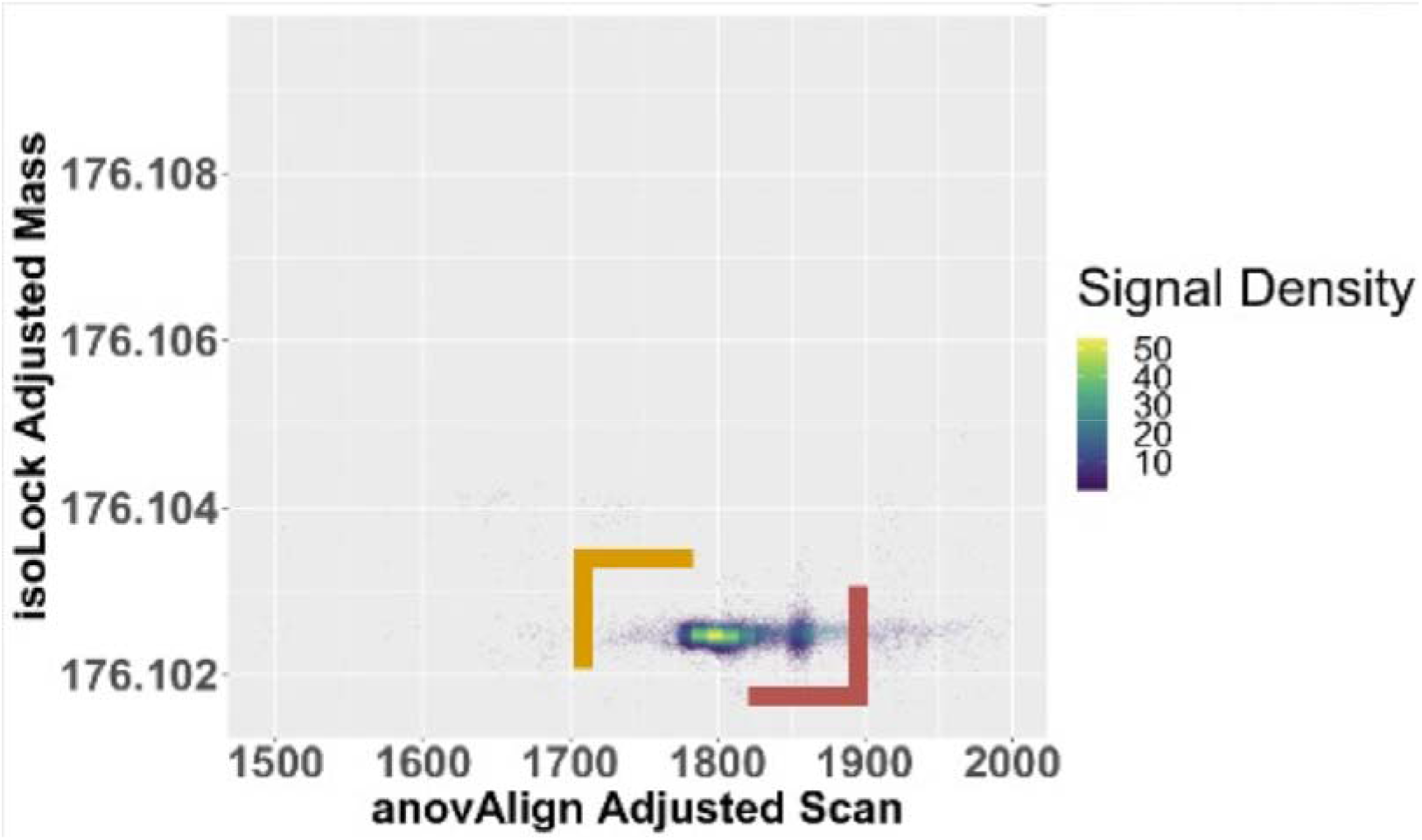
After running our full pipeline and examining the predicted bounds around single returned citrulline mass feature, it is clear that mass drift is corrected by isoLock, the mass is correctly called by autoCredential, and the retention time bounds are accurately called using anovAlign (correctly identifying the citrulline signal as a single mass feature).

We present analyses of two datasets that reflect the ability of our software to capture biologically important metabolite signals. The first dataset is a re-analysis of a previously published untargeted metaoblomics study consisting of hundreds of human microbiome samples. Using this dataset, we show that the suite of isoLock, autoCredential and anovAlign detects the majority of mass features found through a commercial program, Progenesis QI. We also show that a number of mass features, validated via isotopologue signals, are uniquely detected using our software (Figure 11). The second dataset is a smaller untargeted plant metabolomics experiment, but contains 4 replicate samples from 3 *Setaria viridis* genotypes under two watering conditions at 3 time points, allowing for the comparison of the percentage of variance accounted for by the experimental factors. Using this second dataset, we show that our pipeline outperforms the open source software, XCMS, in terms of detecting numbers of mass features and those with high levels of variance explained by genotype, time and treatment (Figure 12).

**Figure 11:**
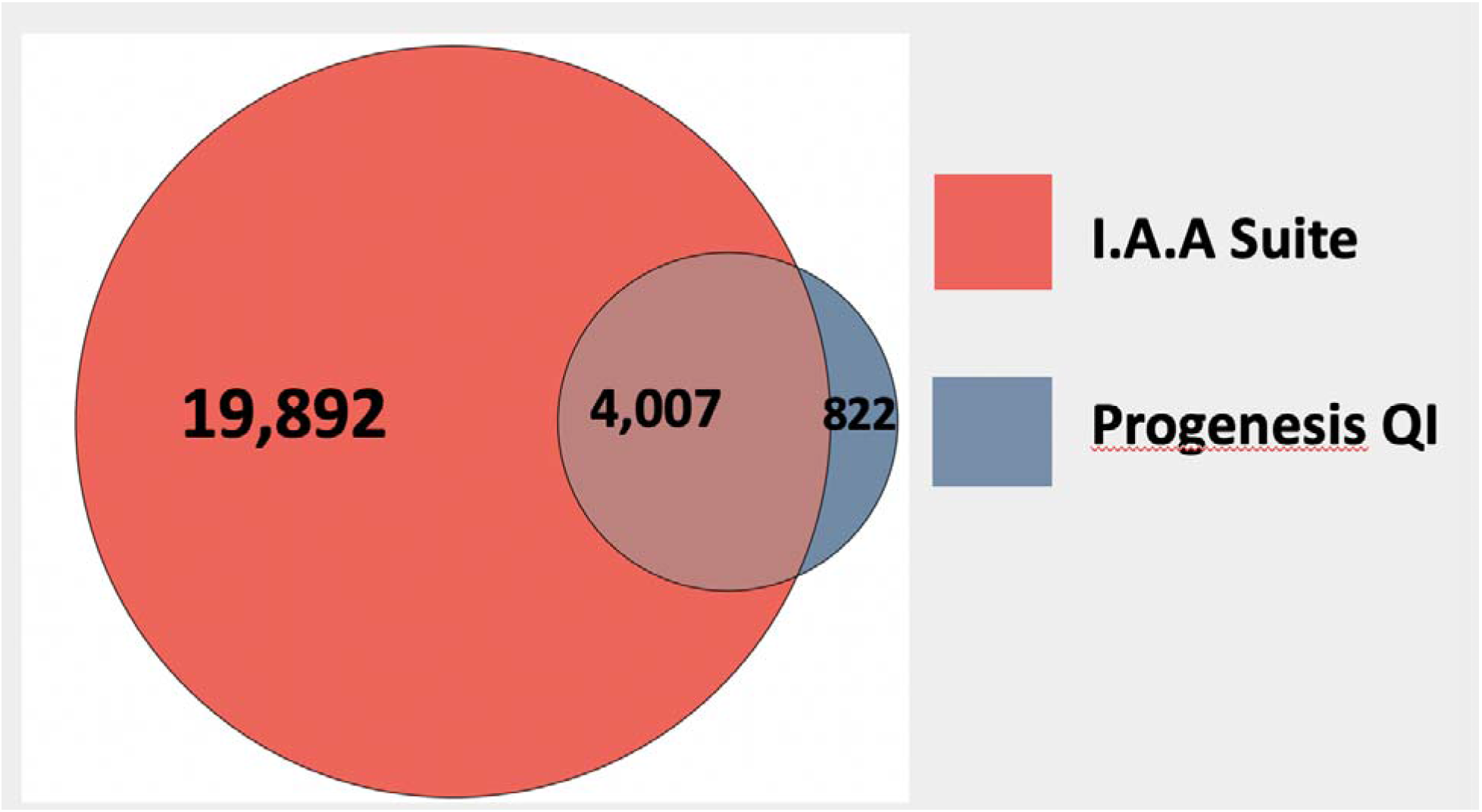
High Priority Mass-Feature Signals at +/- 15 ppm and +/- 3.5 min. Analyzing a large (600 samples) previously published human clinical dataset (Lloyd-Price *et al*., 2019) demonstrates that the isolock, autoCredential and anovAlign suite captures the majority of mass features predicted by a commercial software, Progenesis QI. We also show that our pipeline predicts thousands of additional mass features.

**Figure 12:**
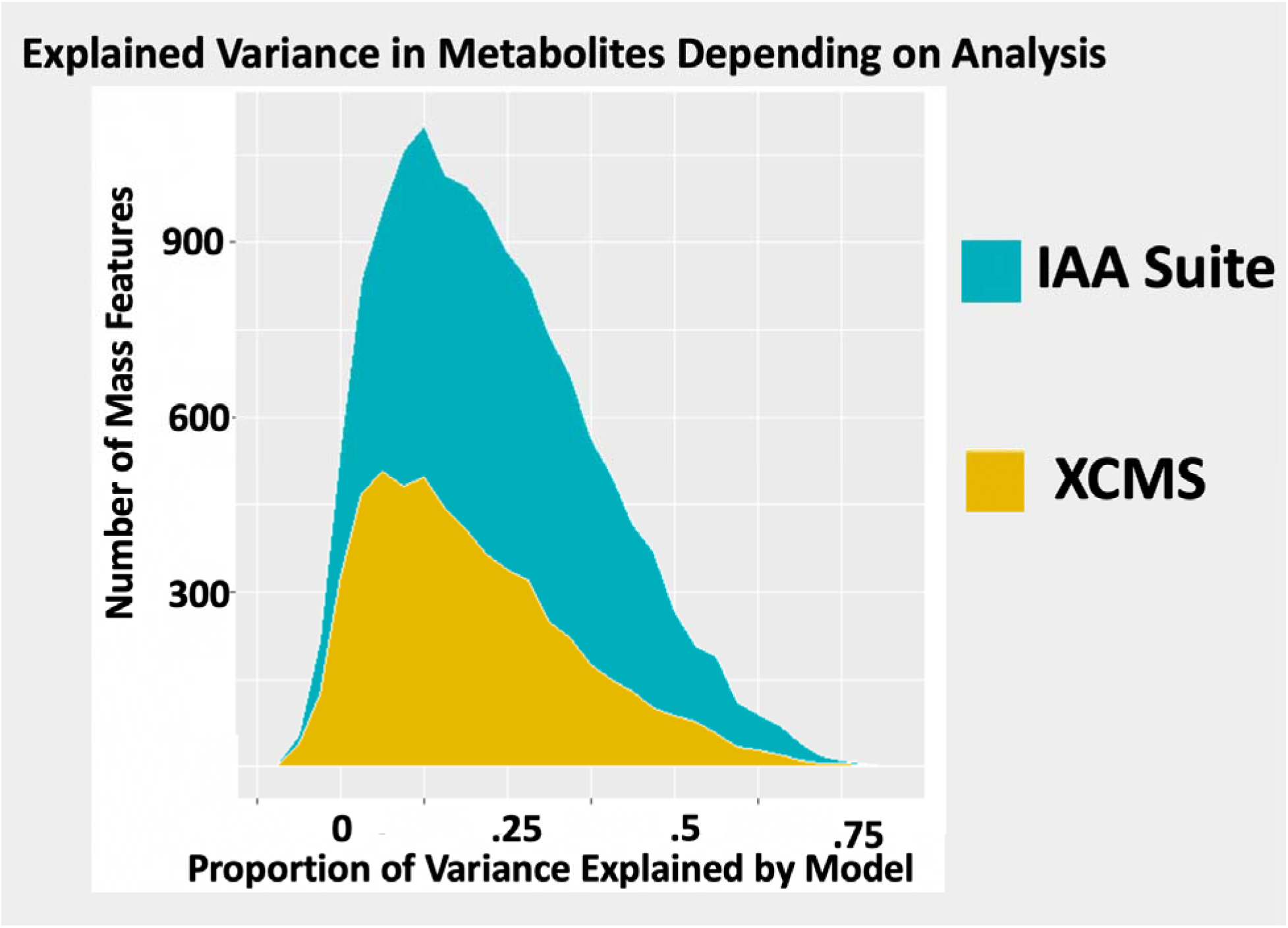
The IAA suite outperforms XCMS. Using a plant dataset with replicates and three experimental factors (genotype, treatment and time) enables the calculation of variance explained by the factors. A. Number of compounds detected plotted against the variance explained by the model (Intensity ~ genotype + time + treatment + genotype*time*treatment).

**Figure 13:**
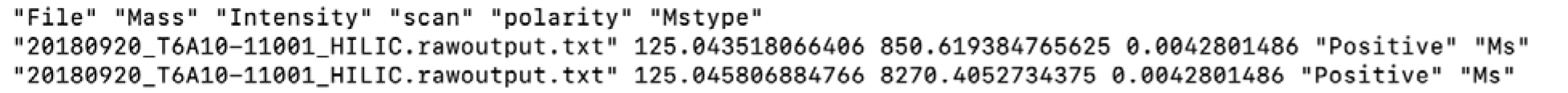
Input to the IAA suite requires .tsv’s with columns containing only data relevant to construction of a merged mass spectra and subsequent peak picking (name of file, Mass, Intensity, retention time, polarity and MsType)

### Application

## Methods

Current peak picking paradigms identify metabolite mass features as 2-dimensional regions of likely metabolite signal known as regions of interest (ROI’s). The first innovation of the IAA suite is to recognize that this problem can be rephrased as a much simpler one: simply taking the intersection of two arrays in order to identify isopairs. Mass spectrometry data are typically reported to multiple decimal points beyond the resolution of a high-resolution mass spectrometry machine. At even single ppm resolution, only the 4^th^ to 5^th^ decimal points are relevant for effective mass resolution within relevant small molecule mass ranges (80-1200 Da). Thus, a complete mass spectra can be effectively discretized by rounding every mass-intensity signal to the fifth decimal point without a loss of resolution. This discretization is critical for determination of important statistical attributes, without incurring information loss. Discretization also enables rapid calculation of isopairs. In order to provide an interface between raw mass spectra and the IAA suite, vendor binary converting libraries (RawFileReader 5.0) are used to convert .raw files to tab delimited text files of the following format: The IAA suite utilizes extensive parallelization to manage the hundreds of gigabytes to terabytes of data contained in a large scale LC-MS study. Parallelization and sequential execution of all steps in the IAA suite (whether accomplished via Python or R) are managed via a Python dask workflow and interface with a robust job engine (such as HTCondor).

To accommodate large-scale calculations, these .txt files are then converted to hdf5 files using the python package Vaex. Hdf5 is a hierarchical data format that allows rapid manipulation of larger-than-memory datafiles. These hdf5 files are manipulated in python (Python 3.7.4) for all subsequent calculations involving isopairs (Figure 14). Isopair calculations on a mass spectra involves two arrays representing the mass values from a spectra containing data from one or more hdf5 files. Isopairs are calculated via the following operations, which are essentially set intersections. Isopairs are fundamental to isoLock and autoCredential.

**Figure 14:**
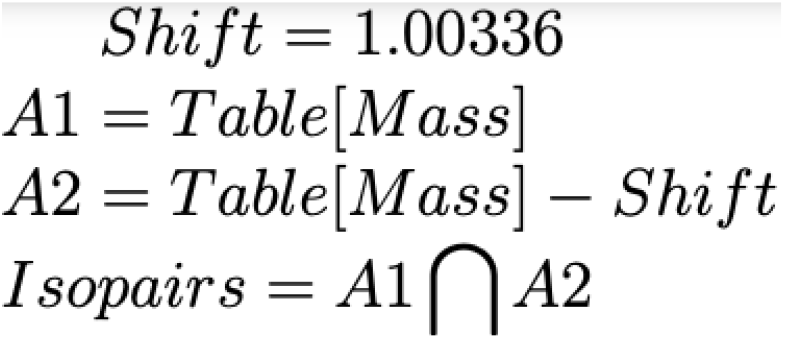
isopairs, which are used extensively in the IAA suite, are calculate by determining signal pairs in raw mass spectra with mass separation equivalent to the mass gain between a 12C an 13C atom (1.00336 Da)

### IsoLock Calculations

During isoLock (Figure 15), isopairs are calculated in an iterative fashion. A1 is the array of masses from a reference file, while the array A2 contains the masses from a query file. In each iterative calculation of isoLock, isopairs are calculated between A1 and A2. In each iteration, the shift between isopairs will vary across a range of 1.00336 (the mass gain of a 13C vs 12C element) +/- plausible drift values between samples (up to 20 ppm). The most likely value of drift is the shift which maximizes the number of isopairs. This drift value is then subtracted from the query file in order to align it with the reference This drift value is then subtracted from the original mass and added as a new column in the hdf5 file, providing a column of drift corrected mass. isoLock can be run on each file in parallel, allowing rapid drift correction of thousands of files. isoLock is implemented in Python, using the Vaex library to manipulate hdf5 files.

**Figure 15:**
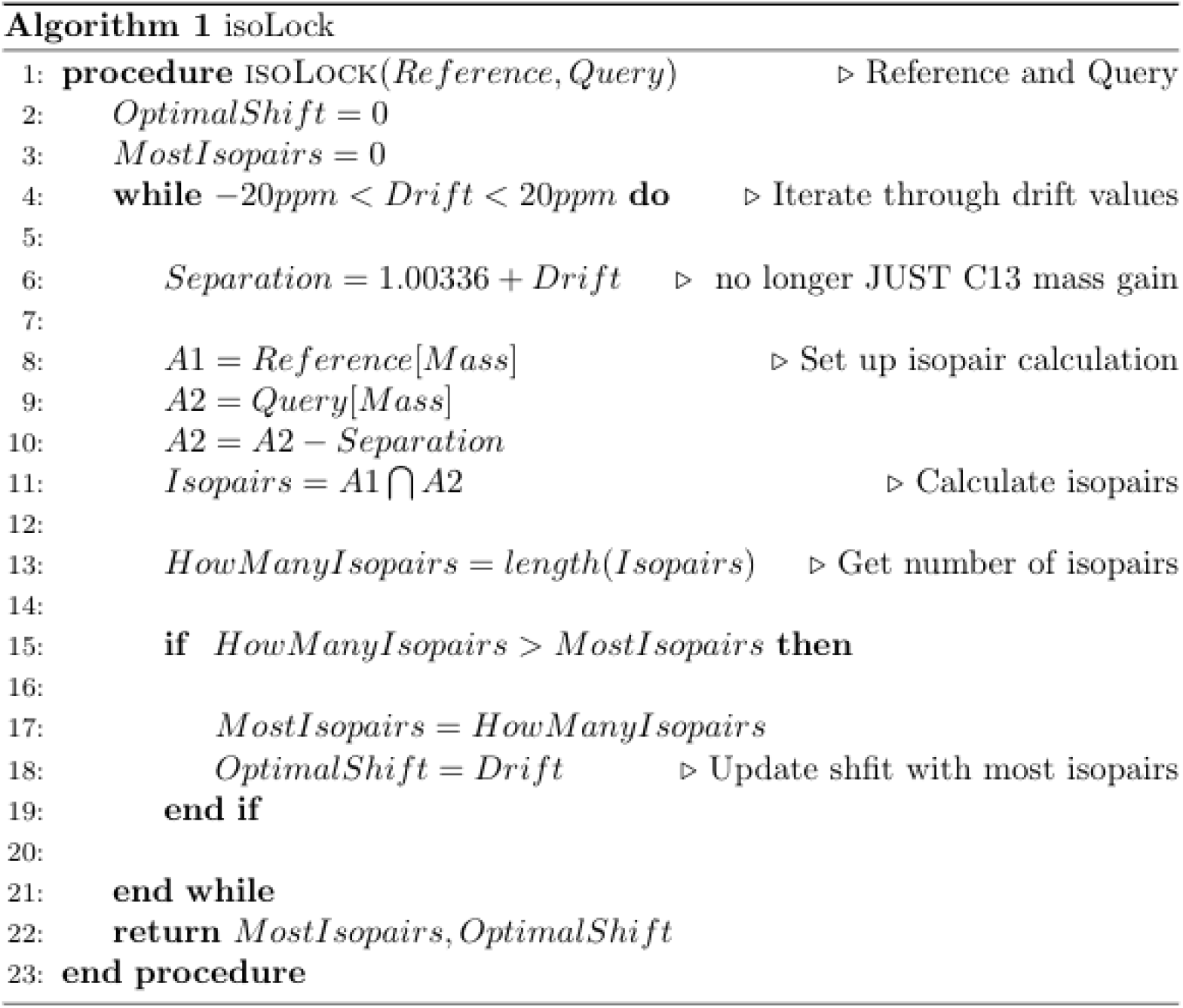
isopairs are used to determine mass drift between a reference and query sample. One array used in the calculation of isopairs(A1) contains the mass signals in the reference sample, while another (A2) contains the mass signals in the query.

### autoCredential

autoCredential (Figure 16) uses files isoLock corrected files in to create a merged mass spectra. Isopairs are used to identify metabolite masses from the merged mass spectra. The merged mass spectra object is created and manipulated in Python via the Vaex and Pandas libraries. In addition to containing the M0 and M1 signals needed to calculate isopairs, the merged mass spectra also contains noise regions. However, using the enhanced signal to noise qualities of a merged mass spectra, these are easily identified and removed via denoising. This is accomplished via permutation testing in R. Resampling of randomly generated masses determines the median signal count in .0001 Da mass bin in noise regions. Bins of the same span (.0001 Da) which do not surpass this threshold are removed. From the remaining space of signals, isopairs are calculated, thereby determining the small fraction of masses associated with carbon containing mass-features. Because single metabolite may create multiple isopairs, masses within 1ppm are collapsed (the .00001 Da resolution of each isopair is within this 1ppm, range for relevant small molecule masses).

**Figure 16:**
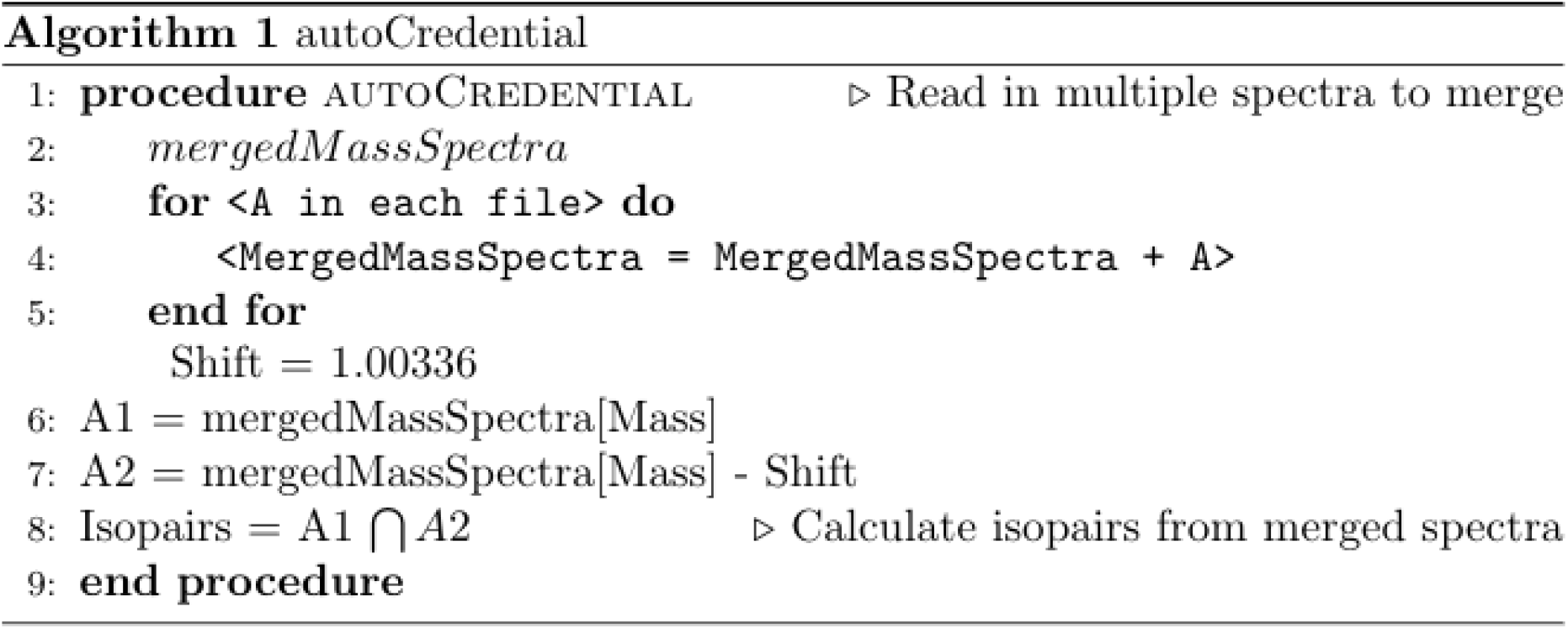
In autoCredential, isoLock adjusted samples are used to create a merged mass spectra, and isopairs are used to identify only mass signals with corresponding M1’s.

### anovAlign

The anovAlign (Figure 17) algorithm models retention time drift on an individual mass-feature by mass-feature basis. Once masses are accurately determined using isoLock and autoCredential, regions of the chromatogram around each mass (+/- 5 ppm) can be extracted from each sample. In the IAA suite, chromatogram slices are extracted mass by mass from each sample’s hdf5 file and concatenated into merged EIC slices. Each merged extracted ion chromatogram (EIC) slice contains the signal (+/- 5 ppm) associated with a mass across ALL samples. The extraction of these regions is parallelized by file, using the Python package Vaex to query each hdf5 file.

**Figure 17:**
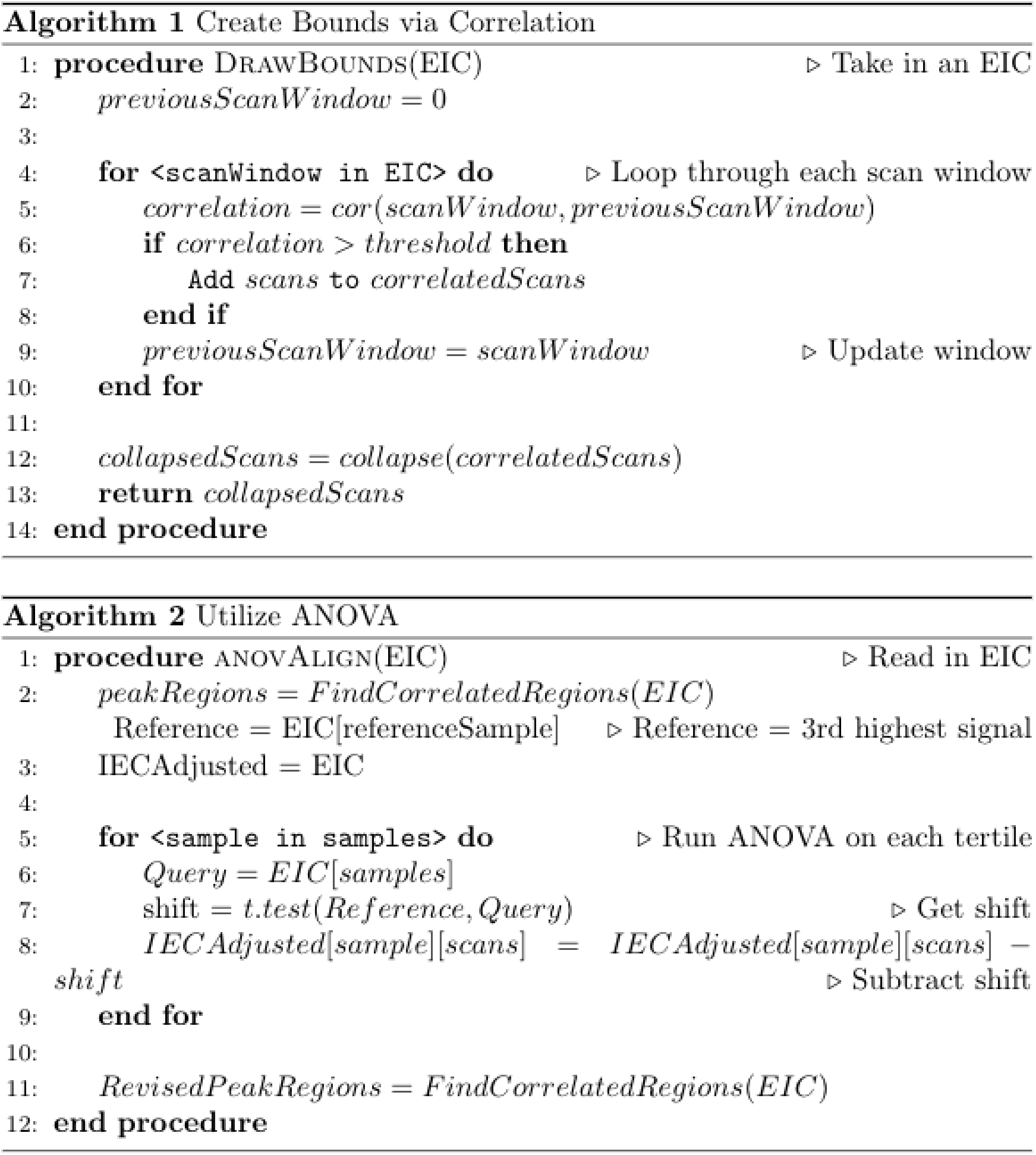
In anovAlign, correlations across local regions of scans are first used to draw approximate boundaries (Algorithm 1). Within these boundaries, ANOVA is used to align signals to a reference (Algorithm 2) before refined bounds are drawn on aligned signals using Algorithm 1.

Modeling drift on a metabolite by metabolite basis in each merged EICslice avoids the common compromises of state of the art warping algorithms such as obiWarp (Prince and Marcotte, 2006) or the more refined warpGroup (Mahieu *et al*., 2016). Warping algorithms, such as obiWarp, attempt to align large regions of chromatogram to one another using global fits. However, the actual drift function for individual metabolites varies substantially across compound classes within these regions. Thus, corrections calculated by current warping algorithms can be biased in favor of certain mass-features, at the expense of others.

anovAlign first denoises a merged EICslice (keeping only the 95^th^ percentile of scans). As an analyte flows through its retention time window, adjacent regions will be statistically similar to each other. This means that adjacent windows will contain data from many of the same files, despite some dropouts, and these signals will correlate with each other. Preliminary bounds (prior to drift correction) are drawn by treating a denoised, merged EIC slice as a grid of 5 scan intervals, and identifying adjacent regions that correlate with one another. Within each of these regions of a potential mass-feature, drift is modeled via ANOVA to produce refined bounds.

In order to implement ANOVA towards alignment, the sample with the third highest signal (to avoid potential outliers) within a region of correlated scans is chosen as an alignment reference. The signal in potential mass feature region is first divided into thirds by intensity. Signals are divided into tertiles for alignment to minimize the effect of intensity dependent drift. Within each tertile of intensity, the average difference in scans between a query sample and the reference is determined via the t.test() function in R (ANOVA). This difference (if significant) is subtracted from the query, in order to align the query signal with the reference. Following the alignment within each provisional mass feature region, final bounds are determined by again searching the merged EIC slice for correlated scan windows, this time using the ANOVA corrected scans. In addition to capturing the intuitive notion of retention time drift as the translation of Gaussians, this approach is not constrained to Gaussian profiles, but merely assumes that drift itself is the introduction of an (approximately) Gaussian source of error. Modeling individual masses in isolation avoids propagation of biases across metabolites, allowing errors to become independent and randomly distributed at a large scale. Structuring the problem in this fashion also sets the stage to easily incorporate standards-based adjustments to anovAlign.

### Data Production and Procurement

#### Broad dataset

600 positive mode HILIC raw Thermo-Fisher QE files were downloaded from metabolomics workbench (project ID: ST000923) from the experiments described in this manuscript (600 human microbiome samples from Lloyd-Price *et al*., 2019). RawFileReader version 5.0 was used to convert binaries to .txt files in order to run the IAA suite. Previously published mass features for the positive mode HILIC quantified from the Progenesis QI were downloaded from the supplementary data at: https://www.nature.com/articles/s41586-019-1237-9. In order to resolve peak splitting in the Progenesis QI, a custom R script was used to collapse mass features within 2 minutes and 5 ppm of one another into a single mass feature, averaging retention time and masses across collapsed mass features.

### Plant Growth Conditions, Harvesting and Data Analysis

As part of a larger water deficit study, 216 Setaria plants representing 3 genoptypes (TB12-48, A10, TB12-201,) were grown at the Donald Danforth Plant Science Center for 7 days prior to transplantation to the Bellwether phenotyper system (Fahlgren *et al*., 2015). Plants were allowed to equilibrate to Bellwether conditions for 6 days before a water deficit treatment (45 percent of field capacity) was implemented on half of the plants. Plants were then harvested at the following timepoints following the equilibration period: 4 Days, 6 Days and 8 Days. Leaves from 3 plants of the same genotype, treatment and timepoint were pooled into a single test tube upon harvesting in order to average out individual plant characteristics. Thus, the 216 plants across the 3 genotypes, 2 treatments and 3 timepoints resulted in 72 test tubes (3 genotypes *3timepoints*2treatments*4 replicates) * 3 = 216. Plants were taken off the phenotyper and the the youngest fully emerge leaf was removed at the node, placed in a 15ml centrifuge tube with stainless steel ball bearings, and placed in liquid nitrogen, until it could be stored at −80C. Then, a paint shaker was used to grind the samples, keeping them cold with liquid nitrogen. Finally, they were aliquoted into 2 ml tubes, weighed, and submitted to the Donald Danforth Proteomics and Mass Spectrometry Facility (PMSF).

### LC-MS Analysis

Samples were resuspended for RPLC by addition of 50 μL of 30% methanol. For HILIC, samples were resuspended in 50 μL of 80% methanol. For both analyses, plate d with resuspension solvent were sealed with RAPID EPS pierceable sealing mats (BioChromato, Kanakawa, Japan) and shaken at 900 rpm at 10°C for 15 minutes on an Eppendorf Thermomixer then centrifuged 2 minutes at 3800 xg to collect the dissolved metabolites at the bottom of the wells and pellet any remaining particulates before transferring 45 μL of each sample to a new well-plate, sealed and stored in a 4°C cold room until use (1-2 days). Just prior to analysis, an additional 15 μL of methanol was added to each well.

LC-MS was performed using a custom built 2D microLC Ultra (Eksigent Technologies, Dublin, CA) attached to a Q-Exactive mass spectrometer (Thermo-Fisher Scientific, San Jose, CA) using electrospray ionization. Data were acquired in either polarity switching full MS only or in data-dependent MS/MS acquisition mode. Full MS scans were taken at a resolution of 70,000 (FWHM at *m/z* 200) with and automatic gain control setting of 500,000 charges and a maximum inject time of 100 ms. MS/MS scans were taken at a resolution of 17,500 (FWHM at *m/z* 200) with and automatic gain control setting of 50,000 charges and a maximum inject time of 50 ms. The top 12 precursors from the previous full M scan were selected with an isolation window of 2 and fragmented with stepped collisional energy of 15, 25 and 35 NCE. Positive ion mode scans were taken with a spray voltage of 4.2 kV while negative ion mode scans used a spray voltage of 3.9 kV. The sheath gas flow, aux gas flow, aux gas temperature and capillary temperature settings were the same for all scans at 15 units, 5 units, 50°C and 250°C, respectively.

For HILIC analysis a custom packed (Higgins Analytical, Mountain View, CA) 0.5 x 150 mm Zic-pHILIC (Merck-Sequant, Darmstadt, Germany) column with 5 μm particle size was used. The solvents used were 10 mM ammonium bicarbonate in water (A) and 10 mM ammonium bicarbonate in 95% acetonitrile. The gradients began with a hold at 100% B for two minutes followed by a linear ramp to 85% B over one minute, then a linear ramp to 50% B over 13 minutes, followed by a linear ramp to 30% B with a hold for one minute before ramping back to 100% B over two minutes with a re-equilibration time of 10 minutes. For RPLC analysis a 0.5 x150 mm TARGA C18 column with 3 μm particle size was used (Higgins Analytical, Mountain View, CA). The solvents for RPLC were 0.1% formic acid in water (A) and 0.1% formic acid in acetonitrile (B). The gradient consisted of an initial hold at 2% B for 3 minutes followed by a linear ramp to 100%B over 10 minutes and a hold for 3 minutes before ramping back to initial conditions over 3 minutes with an 11-minute re-equilibration time. Both methods used a flow rate of 15 μL/minute and injection volume of 2 μL.

### Data Analysis

.raw files were converted to .txt files using RawFileReader version 5.0 in order to run the IAA suite. Thermo Fisher .raw files were converted to mzML format via msConvert (ProteoWizard release: 3.0.20287), downloaded from: https://hub.docker.com/r/chambm/pwiz-skyline-i-agree-to-the-vendor-licenses. Relevant msConvert flags were: --filter “peakPicking true 1-1” --filter “polarity positive” --filter “threshold count .00001 least-intense.” XCMS commands in relevant functions used to process mzML files for peak picking from mzML files were: xcmsSet(method=“centWave”, peakwidth=c(22.25,109.5), mzdiff= .0084, snthresh=5.7, ppm = 3, noise = 100, bw = 6, minfrac = .8), xset2 <-group(xset, bw = 6, minfrac = .8), and retcor(xset2, method=“loess”).

## Discussion

As large scale LC-MS datasets proliferate, robust peak-picking software are needed to reliably convert the gigabytes to terabytes of raw spectra data to mass features. Optimally, these informatics solutions must be sensitive enough to detect biological signals, but have safeguards to prevent flagging non-metabolite signals (salts, contaminants) as mass features. They must also contend with drift in the mass and retention time domains. Our software suite addresses the common problem of “noisy data in, noisy-data out” issue encountered by many LC-MS pipelines by first leveraging isoLock to correct for mass drift. This represents a significant improvement in the ability to quantify mass features in LC-MS data. Essentially, this software suite provides a statistical workaround to the fact that metabolomics lags behind the other -omics partly because the discipline lacks a foundational technology equivalent to polymerase chain reaction (PCR). While other -omics (i.e. many forms of genomics and transcriptomics) frequently overcome signal to noise issues by using PCR to physically amplify genetic material, no such resource exists to selectively amplify metabolites. We demonstrate that this should no longer be a stumbling block, however, as merging mass spectra and chromatograms can exponentially increase signal to noise ratios, and paves the way for new, more powerful peak-picking algorithms. Our suite addresses the concerns that arise with pooling data into a merged data file such as mass drift (isoLock) and retention time drift (anovAlign).

Application of the IAA suite to real world data demonstrates that it improves sensitivity, without incurring the tradeoff of false positives. This is accomplished, generally, by formulating the computational problems of LC-MS peak picking in a way such that the law of large numbers is utilized advantageously. For example, while individual spectra are noisy, pooling samples increases signal to noise ratios. This is somewhat counterintuitive, but immediately evident, when the data is examined. On average, signal hyperconcentrates around metabolite masses across multiple runs. This pooling would, ordinarily, cause signal smearing due to mass drift. However, solving for mass drift (isoLock) beforehand avoids this complication. This can explain many of the significant advantages of our workflow when re-analyzing the previously published dataset from the Broad Institute. Many of the peaks in this dataset demonstrated substantial splitting (when processed using Progenesis QI) due to noise in both the mass and retention time domains. Resolving these issues via isoLock did not reduce sensitivity, and rigorous denoising via permutation tests, assured restriction of false positives. This made it possible for the IAA to recapitulate the vast majority of high priority peaks predicted by Progenesis, while being able to detect many more.

The second application of the IAA suite, on a smaller but more complex plant metabolomics dataset (multiple biological replicates across genotype, time and treatment) demonstrates that these computational advantages translate into enhanced biological signal, compared to peaks quantified using a typical XCMS workflow. This analysis shows that the IAA suite results in not only more peaks but also, peaks with greater average variance explained by a biological model.

Importantly, the principles underneath these algorithms are highly generalizable and our suite is modular – allowing effective application to other workflows besides LC-MS, such as direct injection experiments which are popular in burgeoning single cell metabolomics, a context which could benefit immediately from these tools. These tools will also have utility to other contexts which require accurate LC-MS signal processing beyond untargeted metabolomics, such as proteomics or the use of mass spectrometry for non-traditional analytical settings, such as satellite applications in the space sciences (Arevalo *et al*., 2020). Additionally, autoCredential can be modified such that the mass spacing in an isopair is reflective of the mass gain of various chemical species (not simply the difference between 12C and 13C), making this pipeline amenable to labeling experiments and detection of inorganic compounds.

Despite the improvements represented by our software package, significant hurdles remain in LC-MS informatics. This software provides an excellent platform whose future development can address these issues. One such challenge is to match mass features to compounds of known molecular identity. Another related problem is how to associate adducts and other degeneracies with parent ions. Our suite of isolock, autoCredential and anovAlign provides powerful tools whose future development will address these challenges which are particularly important for pharmaceutical and academic applications. For example, knowing that a predicted mass feature is accompanied by an isotopologue provides confidence that an ion is not a salt or artifact of noise. Future algorithms will leverage the isotopic signature of polished mass spectra to not only determine metabolite masses, but also the most likely number of carbons via modeling the decay in intensity between M0 and M1 peaks. This will improve confidence in the annotation of degenerate mass features, by ensuring that adducts (and higher level isotopologues) agree in the molecular formula of their parent ions, at least in terms of the number of carbons and improve database matching. Future modifications of these algorithms will also be applicable to Ms2 data, further improving the ability to translate signal into biological insight via determination of the molecular formula for high confidence peaks. Ultimately, we believe that the extremely effective peak picking capacities of our software suite (isoLock, autoCredential and anovAlign) will allow LC-MS driven metabolomics to affordably and accurately interrogate the complete metabolome.

## Software availability

Code for each of the algorithms is available at https://github.com/kilgain/MassSpec/tree/main.

## Acknowldegements

Special thanks to Noah Fahlgren and Josh Rothaupt for providing informatics services as part of the Donald Danforth Plant Science Center Data Science Core. We would also like to thank the team behind the previously published data cited in this manuscript (Lloyd-Price *et al*., 2019) for making their dataset publicly available, providing an invaluable public resource in the form of a large and high-quality dataset..

AH, SK and BH have filed an invention disclosure related to this work

## Supplementary Figures

**Supplementary Figure 1:**
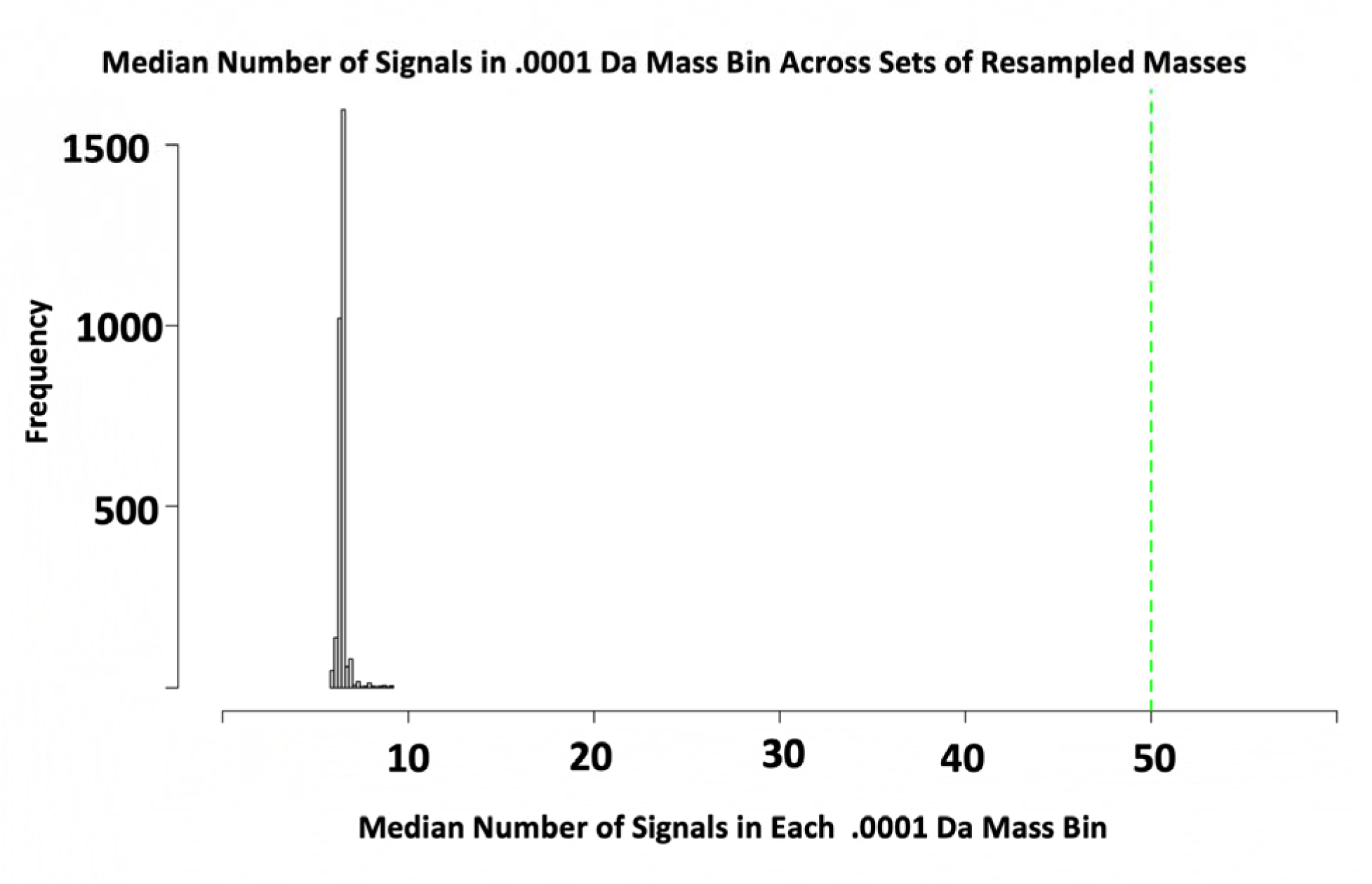
1,000 Masses were chosen at random. A mass slice of 5 ppm around the region of each mass was then selected in order to create a signal profile representatitve of the region around the mass. These mass regions were then randomly resampled into subsets of 100 masses, thus creating a representative subpopulations of mass-intensity background signals. The median number of signals in each .0001 Da Bin in each 100 Mass subpopulation was then calculated in order to determine the number of signals expected in each .0001 Da Mass region due to purely background noise signals. The cutoff for autoCredential (50 counts per each .0001 Da)

